# Artemisinin-independent inhibitory activity of *Artemisia* sp. infusions against different *Plasmodium* stages including relapse-causing hypnozoites

**DOI:** 10.1101/2021.08.10.455849

**Authors:** Kutub Ashraf, Shahin Tajeri, Christophe-Sébastien Arnold, Nadia Amanzougaghene, Jean-François Franetich, Amélie Vantaux, Valérie Soulard, Mallaury Bordessoulles, Guillaume Cazals, Teun Bousema, Geert-Jan van Gemert, Roger Le Grand, Nathalie Dereuddre-Bosquet, Jean-Christophe Barale, Benoit Witkowski, Georges Snounou, Romain Duval, Cyrille Y. Botté, Dominique Mazier

## Abstract

Artemisinin-based combination therapies (ACT) are the frontline treatments against malaria worldwide. Recently the use of traditional infusions from *Artemisia annua* (from which artemisinin is obtained) or *A. afra* (lacking artemisinin) has been controversially advocated. Such unregulated plant-based remedies are strongly discouraged as they might constitute sub-optimal therapies and promote drug resistance. Here, we conducted the first comparative study of the anti-malarial effects of both plant infusions *in vitro* against the asexual erythrocytic stages of *P. falciparum* and the pre-erythrocytic (i. e., liver) stages of various *Plasmodium* species. Low concentrations of either infusion accounted for significant inhibitory activities across every parasite species and stage studied. We show that these antiplasmodial effects were essentially artemisinin-independent and were additionally monitored by observations of the parasite apicoplast and mitochondrion. In particular, the infusions significantly incapacitated sporozoites, and for *P. vivax* and *P. cynomolgi,* disrupted the hypnozoites. This provides the first indication that compounds other than 8-aminoquinolines could be effective antimalarials against relapsing parasites. These observations advocate for further screening to uncover urgently needed novel antimalarial lead compounds.

## Introduction

Sustainable control and elimination of malaria mainly relies on effective treatment, as well as the ability to eliminate the dormant forms (hypnozoites) responsible for the relapses of the widely distributed *P. vivax*^1^. Artemisinin-based combination therapies (ACT) are the frontline treatments against malaria worldwide ^2^. The recent emergence and spread of ACT-resistant *P. falciparum* strains in South-East Asia, is of major concern, and a similar emergence in Africa will have catastrophic consequences^3–5^. Elimination of hypnozoites, the dormant hepatic forms responsible for *P. vivax* relapses, can only be achieved at present using 8-aminoquinoline drugs (primaquine or tafenoquine), but their use is restricted by their deleterious effects in persons with glucose-6-phophate dehydrogenase deficiency^1^. Thus, novel antimalarials are urgently needed.

Artemisinin present in *Artemisia annua* (sweet wormwood) is considered to be solely responsible for the plant’s potent activity against the parasite’s blood-stages^6,7^. Recently decoctions of *Artemisia* dried leaves have been controversially advocated as cheaper traditional plant-based treatments for malaria. Such self-administered and unregulated treatments, due to, are strongly discouraged variable and potentially sub-optimal artemisinin content not least as they would promote the emergence of drug resistance^8^. It has been suggested that the efficacy of such remedies also rests with additional plant compounds that can synergise to enhance parasite killing. This claim is supported by the purported anti-malarial traditional herbal teas based on the African wormwood *A. afra*, a species devoid of artemisinin^9^. We have exploited this difference to explore this issue through comparative and extensive *in vitro* assessment of the inhibitory activity of infusions from both plants on *Plasmodium*.

## Results and discussion

Exposure of *P. falciparum* rings (double synchronized with 5 % sorbitol at 8h interval) to infusions from either plant for 72 h inhibited their growth in a dose-dependent manner **(Fig. 1a)**, with that of *A. annua* active at slightly lower doses. The maturation of the exposed parasites appeared to have been affected by the infusions **(Supplementary Data Fig. 1a)**. We opted to examine the biogenesis of the apicoplast and the mitochondrion as surrogates for the viability of the exposed parasites. Thus, detailed morphological examination revealed that apicoplast biogenesis was disrupted in both *A. annua-* or *A. afra*-exposed parasites where apicoplasts failed to elongate and divide at the schizont stage as they normally do in unexposed parasites **(Fig. 1b)**. The parasite’s mitochondria were also affected by both infusions **(Fig. 1c)**. These observations do not imply that the artemisinin-independent inhibitory activity is directly due to an effect on the biogenesis of these organelles. Nonetheless, preliminary data (**Supplementary Data and Supplementary Fig. 2 & Fig. 3**) suggest that this process is affected to some extend for both organelles.

**Figure 1.**
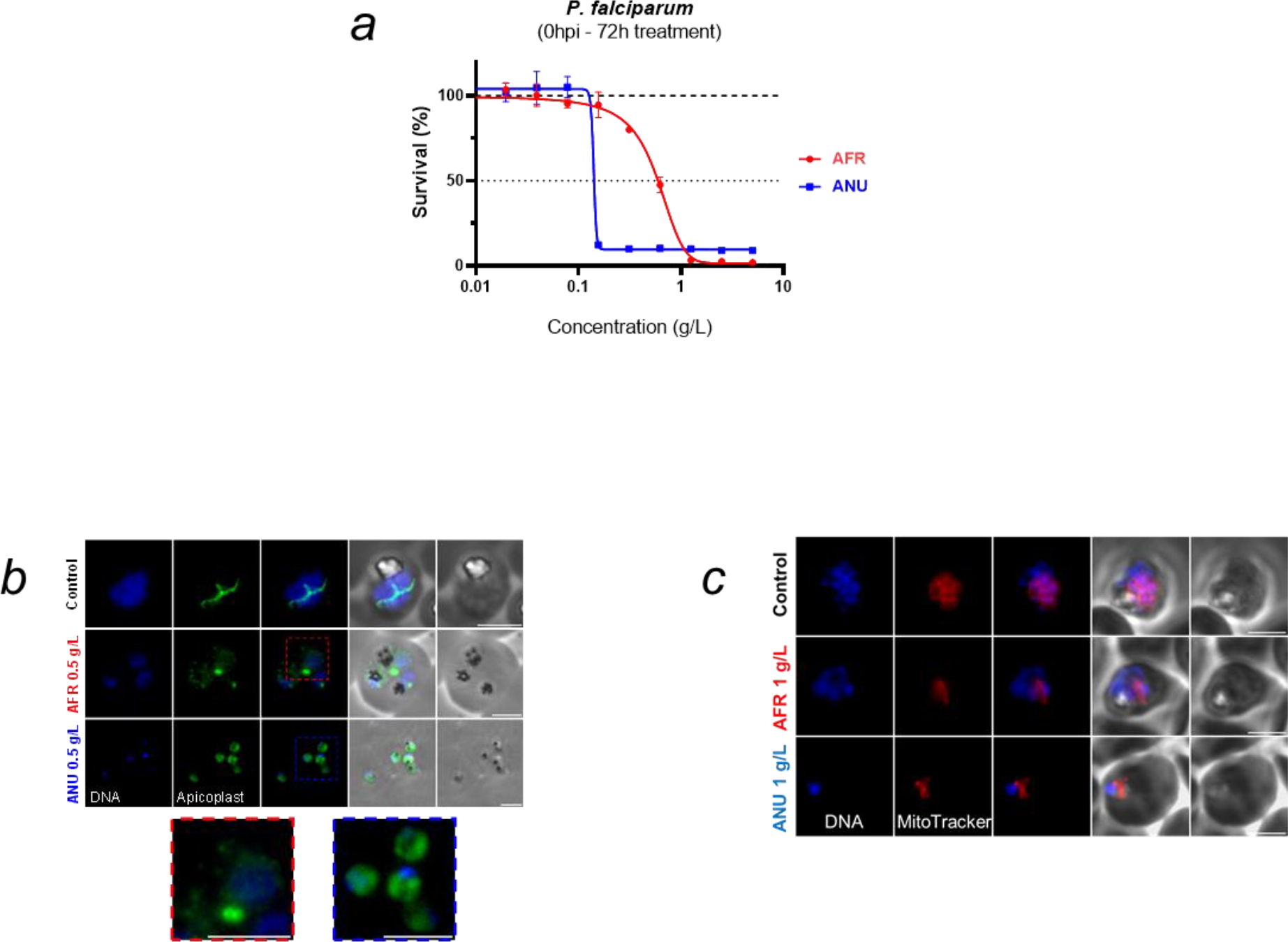
Activity of *Artemisia* infusions on the *P. falciparum* asexual blood stages. **a,** Survival of synchronous ring stage parasites cultivated 72 h in presence of increasing concentrations of *A. afra* (“AFR”) or *A. annua* (“ANU”) infusions. Results are representative of three independent experiments. **b,** Visualisation of apicoplasts by immunofluorescence in untreated control parasites (trophozoite stage) or those grown in the presence of the infusions (*A. afra*, trophozoite stage, red inset; *A. annua*, ring stage, blue inset). An anti-HA antibody was used to localize apicoplasts in a *P. falciparum* cell line expressing a C-terminally HA tagged version of the outer membrane triose phosphate transporter (*Pf*oTPT), a protein known to be located on the outer membrane of the apicoplast^10^. Parasite nuclei were detected by Hoechst 33342 staining. Scale bar is 3 µm. **c**, Visualisation of parasite mitochondria with Mitotracker^®^ Red in infusion-exposed and control cultures. Concentrations of infusions are provided as dry weight of leaves prepared in water and presented as gram per litre (g/L) (See materials and Methods).

We then assessed the infusions against the pre-erythrocytic (hepatic) stages of diverse *Plasmodium* species, where part of the apicoplast metabolism (e.g., FASII activity) is indispensable^11,12^. Exposure of primary simian hepatocytes to the infusions from the time of *P. berghei* sporozoites inoculation and for a further 48 h (spanning the time for full maturation), led to a significant dose-dependent reduction in the size and number of hepatic schizonts, with that of *A. afra* exerting its inhibitory effect at a slightly lower concentration **(Fig. 2a, 2b)**. Similar observations were obtained with *P. falciparum*-infected human primary hepatocytes **(Fig. 2c, d)**, a parasite with a longer hepatic development (5-6 days). Neither infusion had a cytotoxic effect on the hepatocytes, nor did dihydroartemisinin (DHA, used as negative control) affect hepatic schizont numbers though their size was reduced **(Supplementary Data Fig. 4)**. We further demonstrated that exposure of sporozoites to the infusions for 1 h prior to their addition to the hepatocyte cultures, also led to a significant reduction in the number of hepatic schizonts for both parasite species **(Supplementary Data Fig. 5)**. Nonetheless, the viability of such exposed sporozoites, as assessed by vital staining and by their motility, was not altered **(Supplementary Data Fig. 6)**.

**Figure 2.**
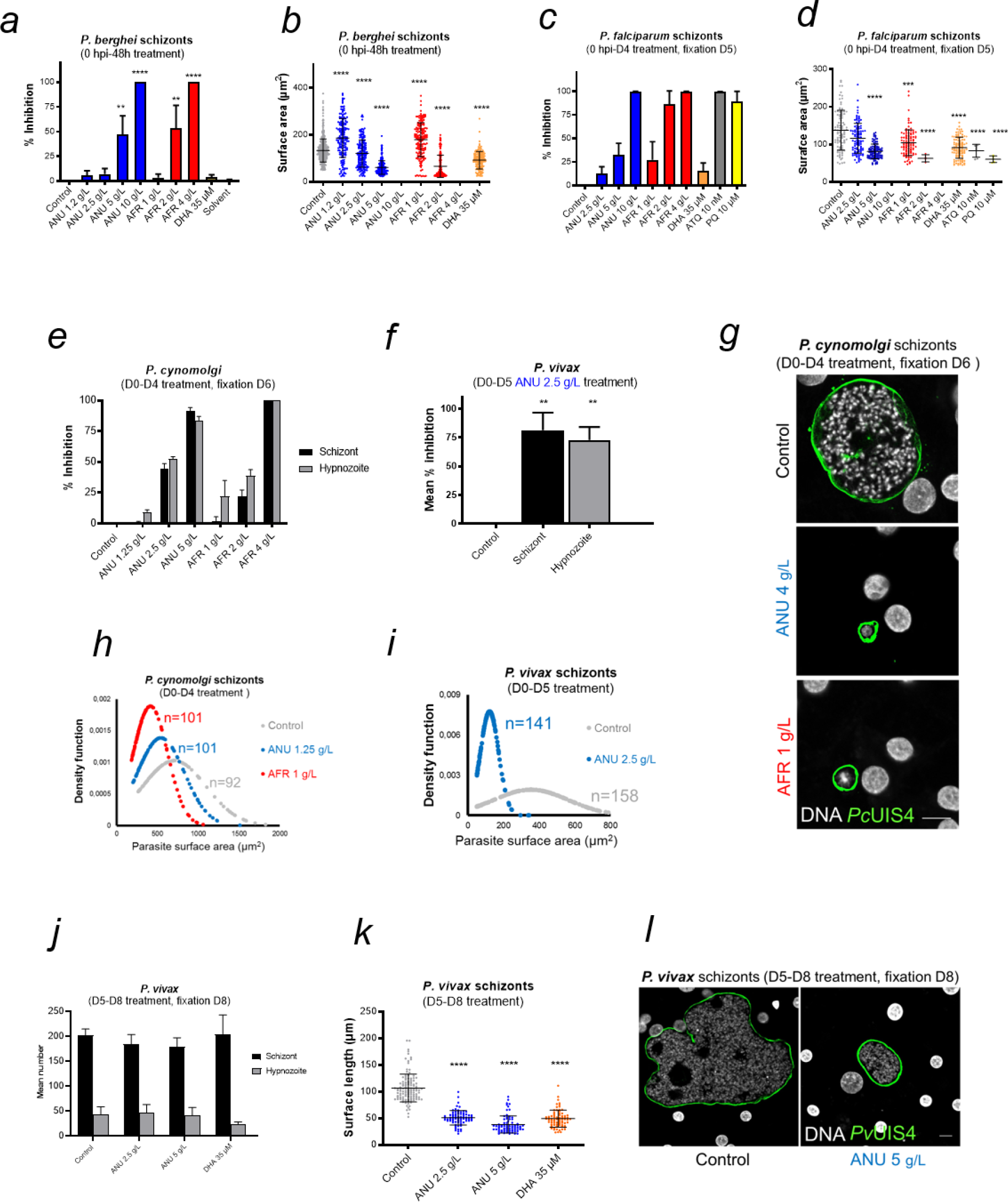
Activity of *Artemisia* infusions on the pre-erythrocytic *Plasmodium* parasites. **a, b,** Quantification of intracellular *P. berghei*-GFP schizont numbers expressed as % of the mean numbers (536 per well) observed in control wells (**a**), and size distribution of these schizonts treated with *Artemisia* infusion and DHA (**b**). 10 g/L *A. annua* and 4 g/L *A. afra* completely cleared parasites from the cultures. **c, d,** Impact of the exposure (D0-D4) of *P. falciparum*-infected human primary hepatocytes on parasite numbers expressed as % inhibition compared to controls (mean of 732 per well) (**c**), and size expressed as surface area (μm^2^) (**d**); 10 g/L *A. annua* and 4 g/L *A. afra* cleared all *P. falciparum* schizonts from the cultures. Results are representative of three independent experiments. **e, f,** Effect of exposure to infusions on the schizont and hypnozoite numbers in primary hepatocytes cultures of *P. cynomolgi* (**e**) or *P. vivax* (**f**) expressed as mean % inhibition. Control cultures had 120 (51 schizonts/69 hypnozoites) and 317 (205 schizonts/112 hypnozoites) hepatic parasites per well of a 96-well plate for *P. cynomolgi* and *P. vivax*, respectively. **g**, Confocal microscopy images of exposed *P. cynomolgi* schizonts. The parasitophorous vacuole of fixed parasites were immunolabelled with anti-*Pc*UIS4 antibody. Host and parasite DNA were visualized by 4′,6-diamidino-2-phenylindole (DAPI) dye (Scale bar = 10 µm). **h, i,** Density function plots of *P. cynomolgi* (**h**) and *P. vivax* (**i**) schizonts size distribution (‘n’ represents the number of schizonts for which the size was measured); the peaks show where most of the population is concentrated. **j, k,** Quantification of *P. vivax* parasite numbers (**j**) and size (**k**) in human primary hepatocytes treated on D5-D8 pi with *A. annua* infusion. **l,** Confocal microscopic images of treated *P. vivax* schizonts fixed at D8 pi, using an anti-*Pv*UIS4 antibody as a parasite PVM marker and DAPI for the DNA marker (scale bar = 10 µm). ANU = *A. annua* and AFR = *A. afra*.

The assays were then extended to two parasite species, *P. vivax* and *P. cynomolgi*^13^ that produce hypnozoites^13–15^. Prophylactic treatment from inoculation and for a further four days (D0-D4 post-inoculation) of *P. cynomolgi*-infected primary simian hepatocytes cultured in the presence of infusions significantly reduced the number of both schizonts and hypnozoites in a dose-dependent manner, by 85% for *A. annua*, and completely for *A. afra* at the highest concentrations **(Fig. 2e)**. Due to restricted number of *P. vivax* sporozoites, only the 2.5 g/L *A. annua* infusion could be tested and this had a slightly higher inhibitory effect on both schizonts and hypnozoites than that on those of *P. cynomolgi* **(Fig. 2f)**. For both *P. vivax* and *P. cynomolgi,* the remaining surviving hepatic schizonts were significantly smaller than those in control cultures **(Fig. 2g-i)**. Although, exposure of *P. vivax* in human hepatocytes on days 5-8 post-infection (D5-D8 pi) to the infusions did not lead to a significant reduction in the number of schizonts or hypnozoites **(Fig. 2j)**, the schizonts showed a significant size reduction **(Fig. 2k, l)** that was similar to that observed in parasites treated with DHA **(Fig. 2j, k)**.

The apicoplasts in the exposed sporozoites were affected in that most of the parasites displayed the dot form and some lacked an apicoplast (“void” category, **Fig. 3a**), in contrast to those in control sporozoites where only 19 % presented as a single dot with the others showing the mature elongated form (probably reflecting the presence in the salivary gland of sporozoites of different ages). For both *A. annua* and *A. afra* infusions, the shift to a high proportion of sporozoites with affected apicoplasts was dose-dependent **(Fig. 3a)**, with no concomitant change to the size, shape, or morphology of the parasite nuclei. The biogenesis of the apicoplast was also disrupted by the infusions in the hepatic stages of *P. falciparum* and *P. cynomolgi*, where an increasing fraction of parasites with “affected” apicoplasts was observed as infusion concentrations increased **(Fig. 3b, 3c)**. For these observations, analysis of the organellar genomes were also consistent with some inhibitory action directed against the apicoplast and the mitochondrion (**Supplementary Data Fig, 2f-h**). The development of apicoplasts in relapsing parasite species has been rarely recorded ^16–18^, and only once for the *P. vivax* putative hypnozoite in a chimeric mouse, where it was found to consist of a single organelle elongating and then branching to form new discrete organelles that appear as a punctate pattern. We observed a similar pattern in the hypnozoites in the *P. cynomolgi* control cultures, while in those exposed to the infusions (D0-D4) we noted apicoplasts with a diffuse staining pattern, implying a disrupted biogenesis **(Fig. 4a)**. Similar infusion-affected hypnozoites were also observed on D10 in *P. vivax* hepatic cultures exposed on D4-D10, though their proportion was less than for *P. cynomolgi* **(Fig. 4b)**.

**Figure 3.**
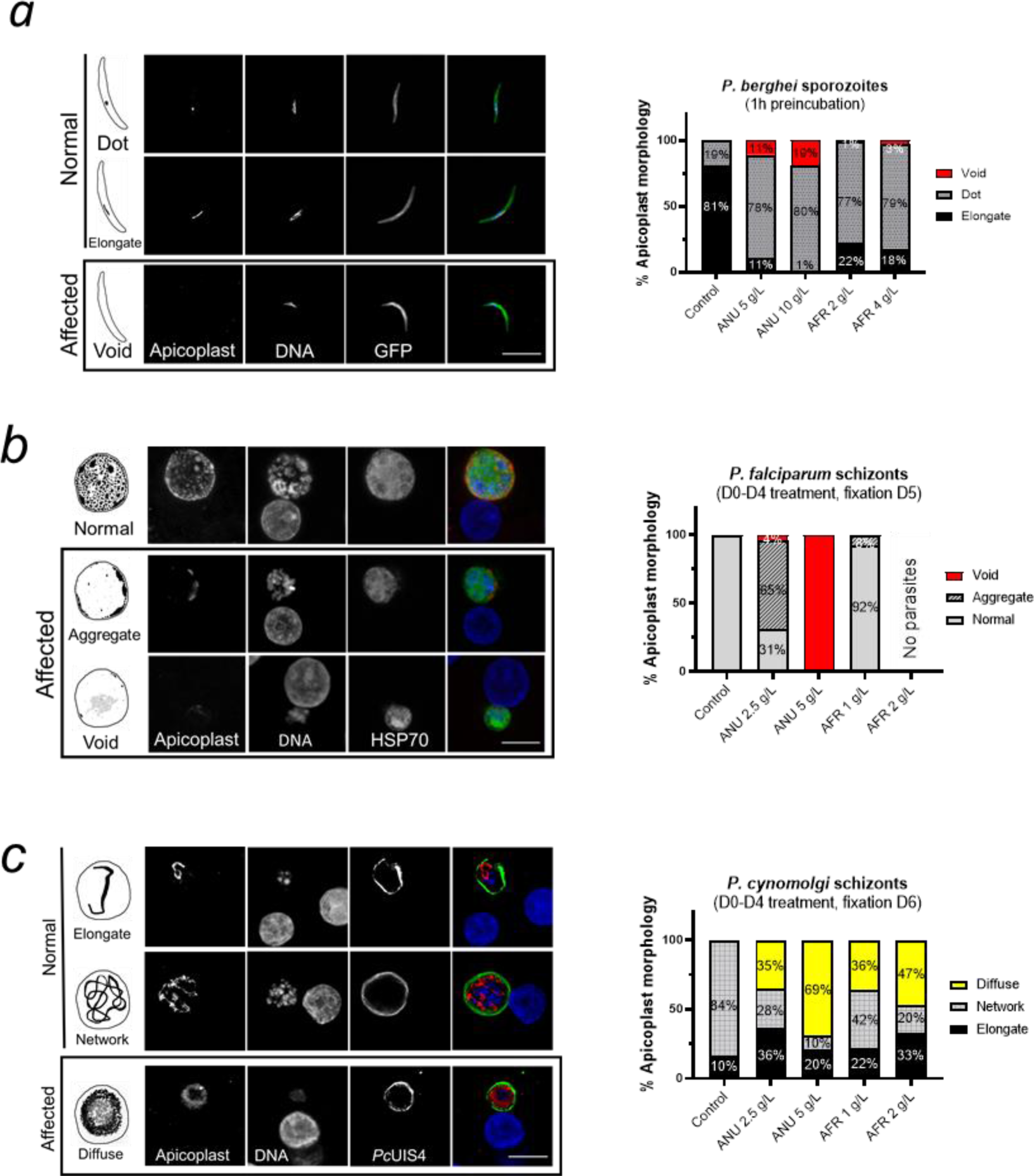
Disruption of apicoplast and mitochondrial integrity in *Plasmodium* pre-erythrocytic stages by *Artemisia* infusions. **a, b, c,** Schematic and confocal images of *P. berghei* sporozoites (**a**), *P. falciparum* schizonts (**b**) and *P. cynomolgi* schizonts (**c**) in control and treated groups. The apicoplast morphologies presented in the “Affected” panels have only been observed in exposed parasites. The frequencies of the various morphologies are presented in the associated graphs. *P. falciparum* and *P. cynomolgi* parasites were detected by anti-HSP70 and *Pc*UIS4 antibodies, respectively. Apicoplasts were labelled by Anti-*Py*ACP antibody (scale bar = 10 μm).

**Figure 4.**
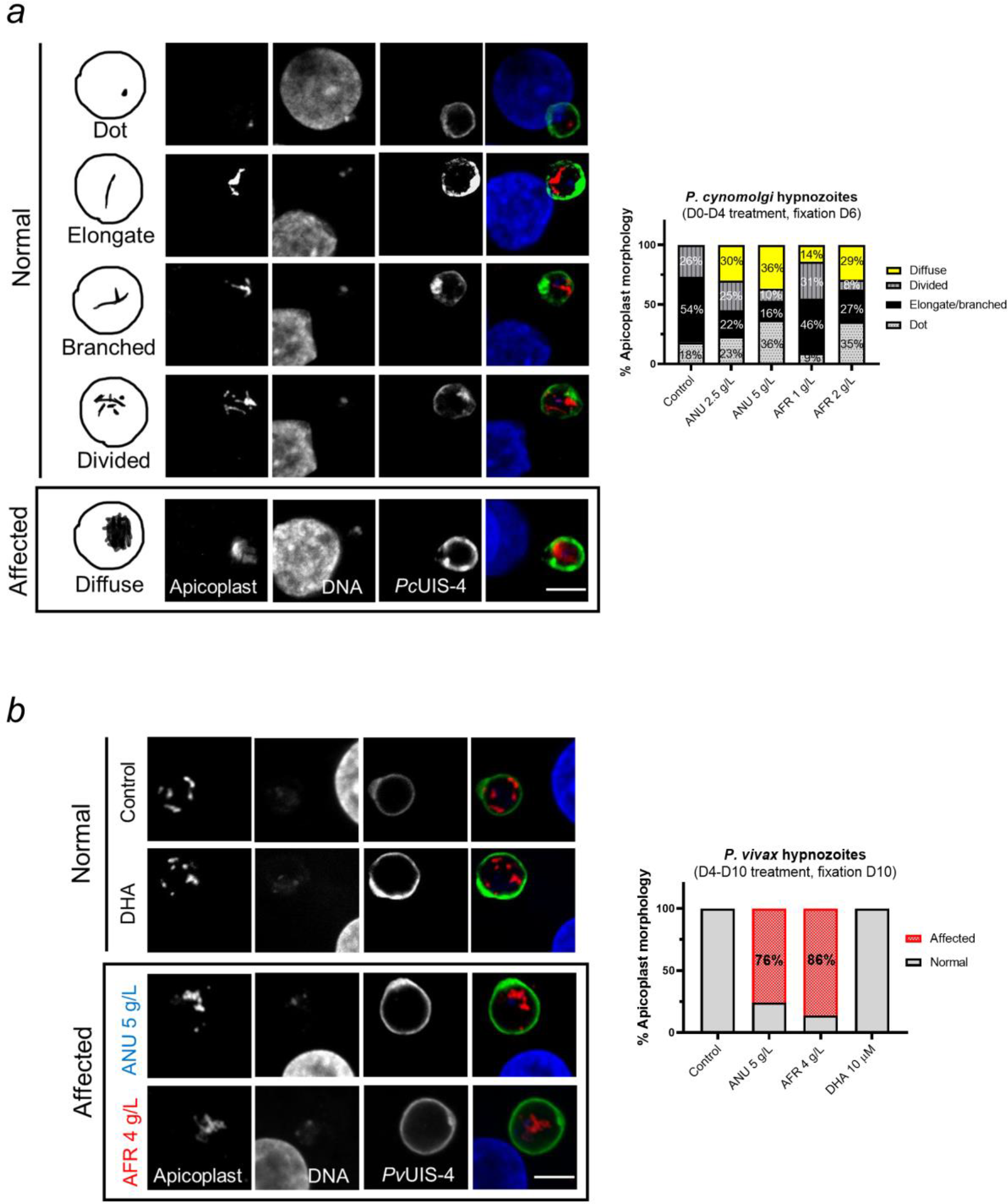
Disruption of hypnozoites’ apicoplast by *Artemisia* infusions. **a,** Schematic (left panels) and confocal images of *P. cynomolgi* hypnozoites on D6 in cultures exposed to infusion during the early hepatic phase. In control cultures, three apicoplast morphologies could be observed, a dot shaped apicoplast in younger hypnozoites that evolves into an elongate organelle that then branches to show a divided morphology as the hypnozoites mature. In infusion-treated cultures (D0-D4), hypnozoites with a diffuse apicoplast signal were additionally observed. **b,** Apicoplast morphology of *P. vivax* hypnozoites exposed to infusions from D4-D10 pi. The apicoplast morphologies presented in the “Affected” panels have only been observed in exposed parasites. Quantification of the various apicoplast morphologies in 100 and 50 parasites for (**a**) and (**b**), respectively, are presented. *P. cynomolgi* and *P. vivax* were visualized by using an anti-*Pc*UIS4 and anti-*Pv*UIS4 antibodies, respectively (scale bar = 5 μm).

The discovery of artemisinin from Chinese traditional medicine has revived interest in *Artemisia* plants as a source of treatments for diverse infectious and non-infectious pathologies beyond than malaria^9,19,20^. The investigations of the *in vitro* anti-malarial efficacy of *Artemisia* infusions have been essentially conducted with the species *A. annua* or of *A. afra*, and this exclusively on the erythrocytic stages (asexual and sexual) of *P. falciparum* parasites ^6,21,22^. Their results are inconsistent and difficult to interpret. Data from our extensive comparative assays clearly demonstrate that infusions from either plant species can fully inhibit the multiplication and the development of both erythrocytic and pre-erythrocytic stages of various *Plasmodium sp. in vitro*. Importantly, this inhibition is independent of their artemisinin content (see **Supplementary Data Table 1** for measurement of artemisinin in both infusions), with maximal activity occurring at broadly similar infusion concentrations, without observable cytotoxic effects against the host cells. Our observations are supported by a recent study investigating the effect of these *Artemisia* infusions on asexual and gametocyte *P. falciparum* erythrocytic stages^7^, that shows an inhibition of the asexual growth after 48 h exposure of both infusions with a significant stronger effect for that of *A. annua*. The higher inhibition levels we observed are probably due to the fact that we assessed parasite survival 72 hours post-exposure (as compared to 48 hours). However, we also observed a strong inhibition effect at 24h and 48h of culture exposed to *A. annua* or *A. afra* 0.5 g/L. Of note, the inhibition of the hepatic stages appears to be mainly due to a detrimental effect on the ability of the sporozoite to achieve a productive hepatocyte infection, which also significantly led to a substantial reduction in hypnozoite numbers in the case of *P. cynomolgi* and *P. vivax*. It is likely that the detrimental effects also observed on the apicoplast in developing *P. vivax* hypnozoite will adversely affect its capacity to reactivate, though this would require further *in vivo* testing. Our *in vitro* observations are clearly insufficient to support claims of *in vivo* prophylactic or curative efficacy for any traditional remedy from the two plants. They do however suggest the presence of hitherto unknown plant components with potent inhibitory activity. Mechanistically, PCR analysis show that some compounds present in the *Artemisia* infusions quickly and directly affect the replication of the parasite’s apicoplast and mitochondria, consistent with some disruption of their biogenesis as observed by immunofluorescence imaging of these organelles in the assays conducted with several parasite species. It thus appears that one or a combination of the myriad of compounds present in the infusion (**Supplementary Data Information**) might target the mitochondrion or the apicoplast directly. The direct correlation between organelle disruption or impaired development and parasite inhibition and death remains to be established.

Ultimately, the relatively similar pan-species and cross-life cycle inhibitory effects resulting from exposure to the two infusions demonstrates the presence of one or more potential lead compounds unrelated to artemisinin. This warrants systematic bio-guided screening to identify them. This is of particular interest, because this might lead to a novel antimalarial class of compound that could effectively and safely prevent *P. vivax* relapses. Indeed, any drugs that could supplant 8-aminoquinolines, the only drugs currently capable of exerting hypnozoitocidal activity^1^, would greatly enhance the prospect of global malaria elimination.

## Materials and methods

### Primary cryopreserved hepatocytes maintenance and infection

In this study, human and simian primary hepatocytes were used. Simian (*Macaca fascicularis*) hepatocytes were isolated from the liver of healthy animals using collagenase perfusion as previously described^23^. Cryopreserved human primary hepatocyte vials were purchased from Biopredic (Lot no. LHuf17905A) or Lonza (Lot no. HUM182641). Cells were seeded in 96-well plates (Falcon by Becton–Dickinson Labware Europe, France) coated with collagen I (BD Bioscience, USA), such that a single cell layer homogenously covers each well. Cryopreserved primary human hepatocytes (Biopredic International and Lonza) were seeded generally 4 days before infection. Both human and simian hepatocytes were maintained at 37 °C in 5% CO_2_ in William’s E medium (Gibco) supplemented with 10% fetal clone III serum (FCS, Hyclone), 1% penicillin–streptomycin (Gibco), 5 × 10^−3^ g/L human insulin (Sigma Aldrich, USA), 5 × 10^−5^ M hydrocortisone (Upjohn Laboratories SERB, France) and Matrigel (Corning, Ref. 354234). The sporozoites used to infect the cultures were purified from dissected salivary glands of infected mosquito^23^. During the infection matrigel cover was removed carefully before adding the sporozoite inoculum. The plates were then centrifuged at 900*g* for 6 min at room temperature before incubation at 37°C for 3 hours. Following a wash, matrigel was added to the culture and allowed to solidify at 37°C for 45 minutes. The cell culture medium was changed thereafter every 24h until cell fixation.

The list of different *Plasmodium* species used for infection of hepatocytes:

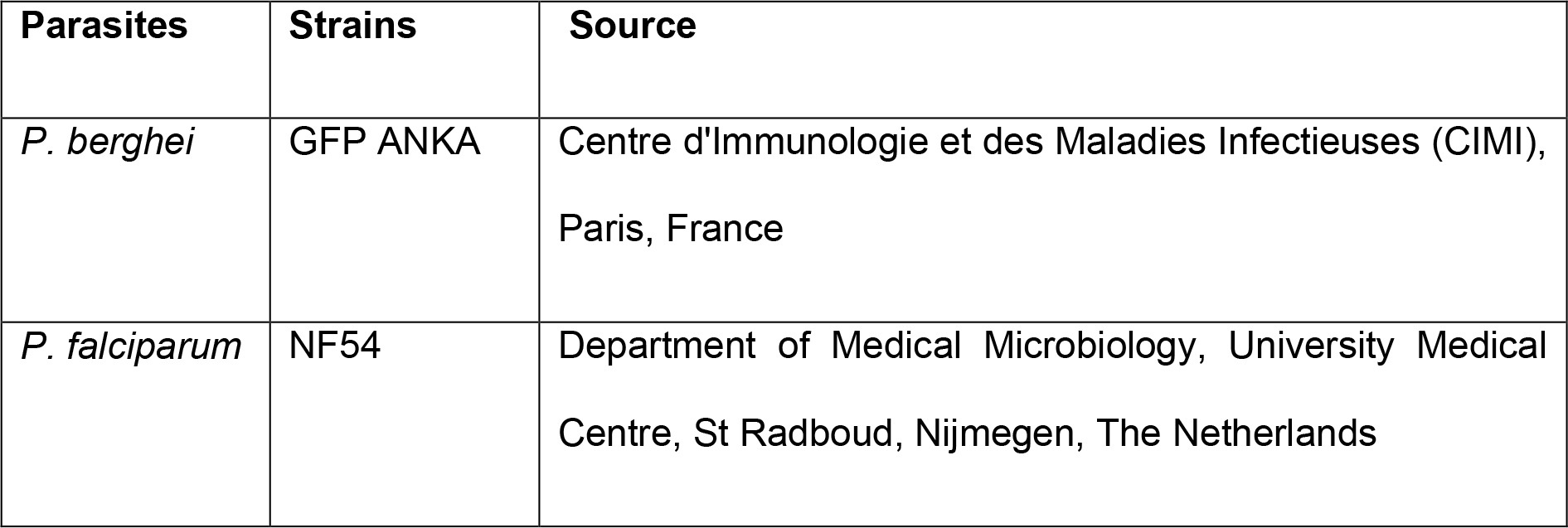
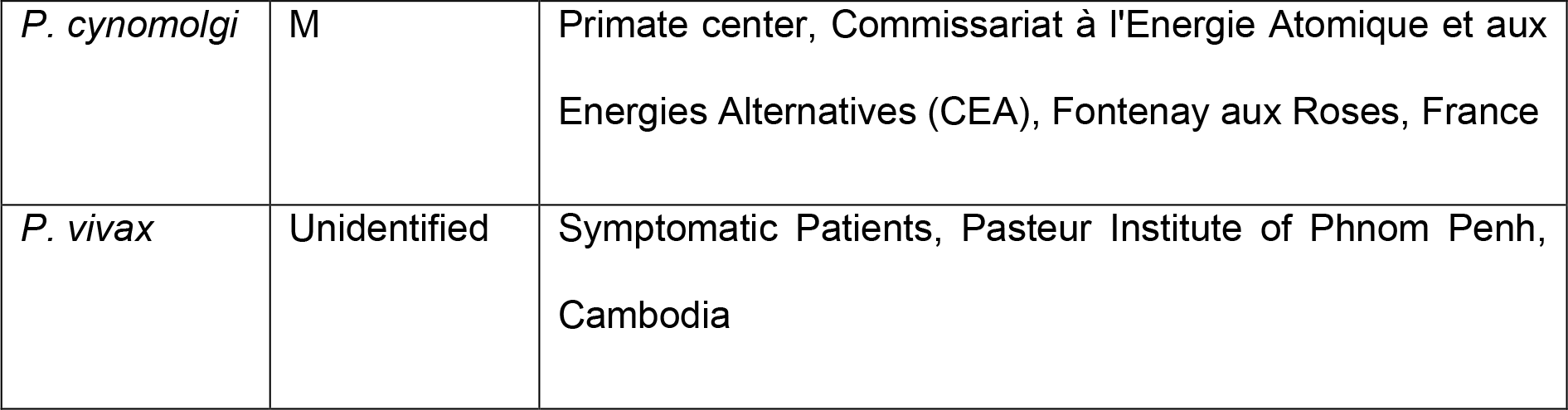

### Culturing *Plasmodium*-infected red blood cells

*Plasmodium falciparum* blood stage parasites were maintained at 2% hematocrit in 1640 RPMI-HEPES supplemented with 5% AlbuMAX II (GIBCO) and 0.25% gentamycin. Parasites were grown in sealed Perspex chambers gassed with beta mix gas (1% O2, 5% CO2 and 94% N2) at 37°C and maintained on 48-hour cycles. Cultures were tightly synchronized at ring stage using sorbitol treatment (%5 v/v) as previously described^24^.

### Production of *P. vivax* sporozoites

Sporozoites were obtained from mosquitoes infected by feeding on symptomatic patients presenting with *P. vivax* in Mondulkiri Province (eastern Cambodia) after obtaining their consent. The patients were managed by the medical staff based at these health facilities. Heparinized blood samples were collected by venepuncture prior to the initiation of treatment, an a diagnostic PCR assay was used to confirm that *P. vivax* was the only *Plasmodium* species present. The blood was centrifuged at 1174*g* at 37°C for 5 minutes, and the plasma was replaced by heat-inactivated naïve human AB serum. Batches of three hundred to six hundred 5-7 days old adult female mosquitoes (*Anopheles dirus*), starved overnight were fed via an artificial membrane attached to a water-jacketed glass feeder maintained at 37°C. Engorged mosquitoes were maintained at 26 °C and 80% relative humidity, and provided with a 10% sucrose plus 0.05% para-amino-benzoic acid solution on cotton pads. Salivary glands of mosquitoes were dissected 14-20 days post-blood meal in L15 medium supplemented with antibiotics and Amphotericin B (Gibco™).

### Production of sporozoites (other species than P. *vivax*)

*Plasmodium falciparum* was transmitted to *Anopheles stephensi* mosquitoes using an artificial membrane feeding on *in vitro* cultivated gametocytes, in Department of Medical Microbiology, Radboud University Medical Center, Nijmegen, The Netherlands. *P. berghei* and *P. cynomolgi* infections of *An. stephensi* mosquitoes were performed in Paris respectively by direct mosquito feeding on anesthetized *Pb*-GFP^25^ infected mice, or by artificial membrane feeding on *P. cynomolgi* (strain M) infected blood from *Macaca fasicularis*. All simian procedures (infection, monitoring and blood sampling) were done in CEA, Fontenay aux Rose before the infected blood samples were transferred to CIMI-Paris for mosquito infection. Two to three weeks post blood meal, *Plasmodium*-infected mosquitoes were killed with 70% ethanol. They were then washed once in Leibovitz’s L-15 medium (Gibco™) containing 5% fetal calf serum, 100 UI Penicillin, 100 µg/ml streptomycin and 0.5 µg/ml amphotericin B, followed by another two washes in the same medium this time lacking serum. The mosquitos were then hand dissected under stereomicroscope and the salivary glands were crushed in a 1.5 ml Eppendorf tube and passed through a 40 µm filter (Cell Strainer, BD biosciences, USA). Sporozoites were finally counted and after dilution adjustment, directly inoculated to the cultures.

### Preparation of *Artemisia* infusion

*Artemisia afra* (PAR, voucher ID: LG0019528 Université de Liège) and *Artemisia annua* (LUX, voucher ID: MNHNL17733 Herbarium Luxembourg) were collected as leaves and twigs and were preserved at room temperature. Both products were protected from sunlight until infusion preparation.

The stock infusion was prepared as follows: 2.5 grams *Artemisia* dried leaves and twigs were added to 25 mL of pre-boiled 25 mL commercial drinkable water (Crystalline Company) and the mixture was then boiled while being stirred for 5 minutes. Following cooling over 10 minutes, the infusion was passed through a 40 µm cell strainer (Falcon, Corning Brand) in order to remove plant debris, and then centrifuged at 3000 rpm for 10 minutes to pellet any remaining fine solids, with a final filtration step over a 0.20 µm membrane filter (CA-Membrane) to obtain a fully clear solution. Filtration was done to eliminate debris produced during plant boiling. During *in vitro* use, debris accumulated on top of hepatocyte culture layer and made parasite quantification difficult. To our knowledge, filtration of *Artemisia* infusions is not performed by patients in endemic areas. The stock infusion (100 g/L) was stored at 4°C (short term storage) or at −20°C (long term storage). For the *in vitro* drug assay, the decoction was diluted to the appropriate concentration (g/L) with William’s E medium (along with supplements).

Frozen and freshly prepared *Artemisia* decoction samples were subjected to chemical analyses including artemisinin content quantification.

### Quantification of artemisinin in infusion: UHPLC Analysis

The chromatographic apparatus consisted of Nexera X2 UHPLC system (Shimadzu, Marne la Vallée, France) equipped with a binary pump, solvent degasser, and thermostatted column compartment. A reversed-phase column was used for separation: Kinetex C18 (2,1 × 100 mm 1,7 μm from Phenomenex). Mobile phase A and B consisted of water (0,1% formic acid) and acetonitrile (0,1% formic acid) respectively. The 10 min linear gradient program used was 10–100% B over 5 min, a plateau at 100% B for 2 min was used to wash the column, decreasing from 100%–10% B in 0,1 min, followed by a 2,9 min post-run isocratic step at 10% B to re-equilibrate the column. The flow rate was constant at 0.5 mL/min at 25 °C.

### Quantification of artemisinin in infusion: MS/MS Detection

MS/MS experiments were carried out using the Shimadzu UHPLC system described above coupled to a Shimadzu LCMS-8050 triple quadrupole mass spectrometer using the multiple reaction monitoring (MRM) technique operating in positive ion mode. The following parameters were used for all experiments: electrospray interface voltage was 3 kV; heat block temperature of 350 °C; desolvation line temperature of 300 °C; interface temperature of 300 °C; drying gas flow gas was set at 5 L/min; and nebulizing gas flow at 3 L/min, heating gas flow rates were set at 15 L/min. Q1 and Q3 were set to unit resolution. The dwell time was set at 1 ms for each MRM transition; optimal CE values were chosen to obtain the most characteristic fragments (see below table).

| Precursor (m/z) | Fragment ion (m/z) | Collision Energy (eV) |
| --- | --- | --- |
| 283.10 | 247.15 | 9 |
| 283.10 | 265.15 | 8 |
| 283.10 | 219.10 | 12 |

### Drugs

Drugs were dissolved in different solvents according to manufacturer’s instructions which are enlisted below: Atovaquone (Sigma-Aldrich, A7986), Primaquine bisphosphate (Sigma Aldrich, 160393), DHA (Sigma-Aldrich, 1200520), Isopentenyl pyrophosphate NH4+ salt (Isoprenoids, LC Tampa, FL 33612), Chloramphenicol (Sigma-Aldrich, C0378-5G), Doxycycline (Sigma-Aldrich, D9891). Media with drugs were changed every 24 h during the assays.

### Immunostaining

Cultured hepatocytes were fixed using 4% paraformaldehyde (PFA) for 10-15 minutes at room temperature. The fixed samples were subjected to immune labelling. The primary antibodies (see below table) were diluted in dilution buffer (1% w/v Bovine Serum Albumin and 0.3% v/v Triton X-100 in PBS) and incubated at room temperature. Primary antibody-stained cultures were washed thrice with PBS 1X and were incubated with secondary antibody (below table) along with DAPI (1:1000) in order to visualize the nuclei at room temperature for 1 hour. Mitotracker® (ThermoFischer Scientific) was used to visualise parasite mitochondria.

List of primary and secondary antibodies used in the study:

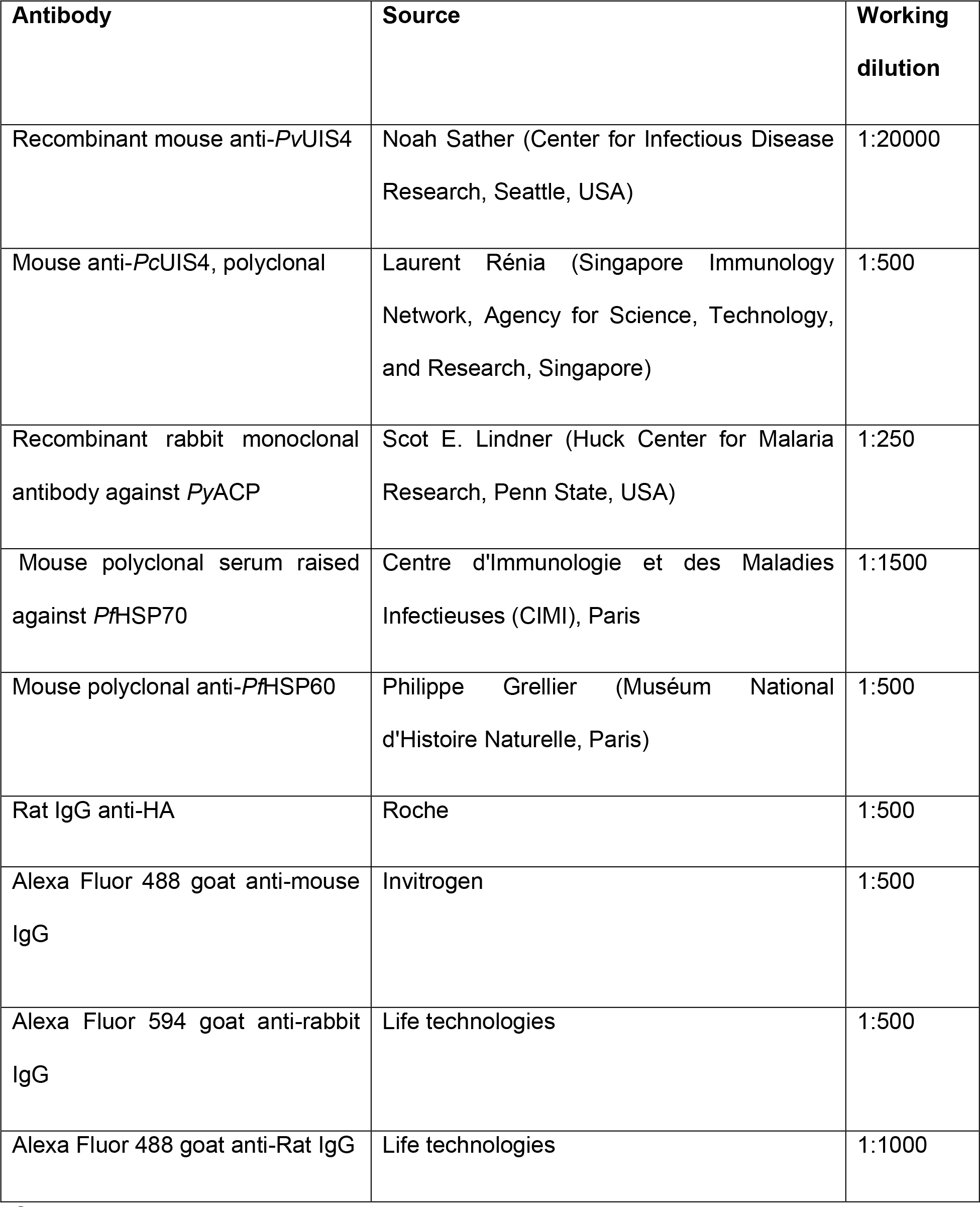

### Confocal Microscopy and Image analysis

Confocal images of immunostained cultures were taken by Leica SP8 white laser microscope controlled by the Leica Application Suite (LAS) AF software at the QUANT microscopy platform (Institut du Cerveau et de la Moelle épinière, ICM) in Paris, France. Z-stack images were constructed and analysed by Fiji package of imageJ software. Final images were made by Microsoft Powerpoint. Schematic figures of *Plasmodium* apicoplast were designed by Procreate® application in Apple ipad pro.

### Quantification of apicoplast morphologies

Pre-erythrocytic and hypnozoite stages: ACP immunolabeled parasites were quantified considering the distinguishable morphology in drug treated versus control by an individual with the aid of confocal (Leica SP8 white laser microscope) at 63X magnification. For each condition, 100 parasites were observed. Erythrocytic stage: TPT-HA immunolabeled parasites were quantified considering the distinguishable morphology in drug treated versus control by an individual with the aid of fluorescence microscope (Axio Imager 2_apotome; ZEISS, ×100 magnification). For each condition, between 54 and 116 parasites were observed.

### Parasite enumeration and toxicity assessment using high-content imaging

Upon fixation and immunostaining, cell culture plates were analyzed in order to determine the number and size of the parasites using a CellInsight High Content Screening platform equipped with the Studio HCS software (Thermo Fisher Scientific). Uninuclear hepatic parasite forms observed beyond D4 were considered to be putative hypnozoites, while multinucleate forms were classed as schizonts. The parasite size reduction was calculated on the average object area using the total surface area of each selected object (µm^2^). The high content imaging approach described previously^26^. To assess cell toxicity of infusion for hepatic cultures, fixed plates were further scanned for the DAPI signal representing host nuclei. The analysis was based on counting of total cell nuclei.

### qPCR to quantify apicoplast and mitochondrion

To perform relative quantification of apicoplast, mitochondrial and nuclear genomes we extracted genomic DNA from *Pb*-GFP infected hepatocytes treated or not with infusion using a NuceolSpin Tissue kit (Macherey-Nagel, Germany) according to manufacturer’s manual. The amount and the purity of the extracted DNA were checked by a Nanodrop ND 1000 machine (Thermo Fischer scientific). The following gene-specific primers were designed to target genes found in each organelle or nuclear genomes:

*tufa*_ PBANKA_API00280 (apicoplast) *5′-GATATTGATTCATCTCCAGAAGAAA-3′* / *5′-ATATCCATTTGTGTAGCACCAATAA-3′*,

Cytb3_PBANKA_MIT01900 (mitochondrion) *5′-AGATATATGCATGCTACTGG -3′* / *5′-TCATTTGTCCCCAAGGTAAAA -3′*

*Green fluorescent protein* (GFP) *5′-GATGGAAGCGTTCAACTAGCAGACC -3′* / *5′-AGCTGTTACAAACTCAAGAAGGACC -3′*,

and *CHT1*_PBANKA_0800500 (nuclear) *5′-AACAAAATAGAGGTGGATTT-3′* / *5′-AATTCCTACACCATCGGCTC-3′*.

The quantification of each target gene was normalized to GFP, expressed as a transgene under the *P. berghei* elongation factor 1α promoter^27^. All reactions were performed on an Applied Biosystems 7300 Real-Time PCR System using the Power SYBR Green PCR Master Mix kit (Applied Biosystems, France), according to the manufacturer’s instructions. The cycling conditions were as follows: initial denaturation at 95°C for 5min, followed by 40 cycles at 94°C for 30s, 56°C for 30s and 72°C for 30s. The experiments were performed in triplicate. Relative quantification of target genes was calculated according to the 2^−ΔΔCt^ method^28^.

### Sporozoite viability assay

Freshly dissected *Plasmodium berghei*-GFP sporozoites were incubated with varying concentrations of *Artemisia* infusion for 1 hour. Then the infusion was removed by centrifugation and sporozoites resuspended in culture medium. One µL of propidium iodide (Gibco) was added to 15000 cells and incubated for 2-3 minutes. A 15 µL aliquot of sporozoite suspension were put on a KOVA^®^ Glasstic microscopic slide (KOVA International) and visualized under a microscope. Finally, dead, and viable sporozoites were quantified under epifluorescence microscope.

### *in vitro* screening assay for sporozoite motility inhibition via video microscopy

Salivary glands (SGs) from infected mosquitoes (17–24 days post infectious blood feeding on Swiss mice infected with *P. berghei*-GFP were isolated by hand dissection and placed into Leibovitz L15 media (Gibco 11415049) supplemented with 20,000 units penicillin-streptomycin (Gibco 15140122) and Amphotericin B. Sporozoites were then released by SG mechanical disruption, filtered through a 40 µm mesh (Cell Strainer, Falcon) to remove SG debris and diluted in activation medium (William’s E medium supplemented with 10% of fetal clone III serum, 1% penicillin–streptomycin, 5 × 10^−3^ g/L human insulin and 5 × 10^−5^ M hydrocortisone) to a final concentration of 400000 sporozoites/ml and kept on ice. Aliquots of 50 µL were inoculated into wells of a 96-well plate (final amount of approximately 20000 sporozoites per well). An equal volume of *Artemisia* infusion dilutions was added to the sporozoites to give a final concentration of (*A. annua 5%, A. annua 10%* and *A. afra* 5%) and mixed by gentle pipetting. The plate was centrifuged for 6 min at 1600 rpm to maximize the sporozoite settlement and immediately placed at RT for 30 minutes. Then the motility of sporozoites was recorded under video microscopy Zeiss Axio Observer 7 at 40X objectives with the GFP excitation/emission filter and pictures were recorded under a frame rate of 1 frame per second with a Hamamatsu Orca Flash 4.0 V3 camera and Zen software for 3 minutes. Notably, the experiments were performed at 37°C. Finally, the tracking of sporozoites were further analyzed with FIJI imageJ and the moving patterns characterized on maximum intensity z-projections. Thereby, sporozoites were classed as gliding if they moved with a circular pattern describing at least one complete circle loop during the 3 minutes acquisition. The percentage residual motile population was then calculated and compared to uninhibited controls (media containing an equivalent amount of water).

### SYBR Green-I-based cell proliferation assay

*Plasmodium* blood stage parasites (regular or apicoplast-free lines) are incubated in 96 well flat bottom plates, 2% hematocrit in 1640 RPMI-HEPES supplemented with 5% AlbuMAX II (GIBCO) and 0.25% gentamycin complemented with appropriate *Artemisia* infusion dilutions and with or with IPP and Cm. Parasites were grown sealed Perspex chambers gassed with beta mix gas (1%O2 5%CO2, 94% N2) at 37°C for 72h. New 96 well black wall flat bottom plates are set up with 100 µL SYBR Green lysis buffer (20 mM Tris, pH 7.5; 5 mM EDTA; 0.008 % (w/v) saponin; 0.08% (v/v) Triton X-100) with freshly added SYBR Green I (1000X), 100 µL from the cultures are transferred and mixed with the SYBR Green lysis buffer and incubated 1 h at room temperature protected from the light. Fluorescence from each well is measured with TECAN infinite M200 plate reader (excitation: 485 nm, emission: 538 nm and integration time: 1000 µs).

### Statistical Analysis

GraphPad Prism 5 (GraphPad. Software, San Diego, CA, USA) and Excel 2016 (Microsoft Office) were used in this study for the data analysis. All graph values are represented by means with standard deviations (s.d.).

## Acknowledgements

Kutub Ashraf received funding from Labex ParaFrap-IRD PhD South program. Shahin Tajeri acknowledges postdoctoral funding support from FRM (Fondation pour la Recherche Médicale) for the PALUKILL project. Nadia Amanzougaghene received postdoctoral salary included in an ANR (L’Agence nationale de la recherche) funded project called Plasmodrug. This work and C.Y.B., and C.S.A. are supported by Agence Nationale de la Recherche, France (grant ANR-12-PDOC-0028, Project Apicolipid), the Atip-Avenir and Finovi programs (CNRS-INSERM-Finovi Atip-Avenir Apicolipid projects), Laboratoire d’Excellence Parafrap, France (grant ANR-11-LABX-0024), LIA-IRP CNRS Program (Apicolipid project), CEFIPRA MERSI program, IDEX Université Grenoble-Alpes and MESRI PhD program fellowship. The investigations on *P. cynomolgi* were funded through a grant from the Agence Nationale de la Recherche, France (ANR-17-CE13-0025-01); IDMIT infrastructure is supported by the French government “Programme d’Investissements d’Avenir” (PIA), under grants ANR-11-INBS-0008 (INBS IDMIT). This work benefited from equipment and services from the CELIS cell culture core facility (Paris Brain Institute), a platform supported through the ANR grants, ANR-10-IAIHU-06 and ANR-11-INBS-0011-NeurATRIS. We would like to thank the help of QUANT microscopy platform of the Paris Brain Institute specially David Akbar, Claire Lovo and Aymeric Millécamps for their help in analysis of microscopic images and sporozoite motility videos. We thank Laurent Rénia, Noah Sather, Philippe Grellier and Scott Lindner for the kind gift of anti-*Plasmodium* antibodies listed in the Methods section. The kind help of Sivchheng Phal and Chansophea Chhin (Institut Pasteur du Cambodge, Phnom Penh, Cambodia) with *P. vivax* experiments is sincerely thanked. We are grateful to Lucile Cornet-Vernet and Pierre Lutgen for providing us *Artemisia* plants.

## Author contributions

K.A., S.T., N.A., J.-F.F., C.S.A., G.C. and A.V. performed experiments.

K.A., S.T., N.A., J.-F.F., D.M., C.S.A. and C.Y.B. analysed data.

K.A., B.W., J.-F.F., C.Y.B., and D.M. designed the study.

J.-F.F., B.W., G.-J.G., G.S., T.B., V.S, J.C.B. and D.M. contributed essential materials.

S.T., C.Y.B., G.S., R.D. and D.M. wrote the manuscript. All authors read and approved the manuscript.

## SUPPLEMENTARY INFORMATION

**Supplementary Data Table 1.**
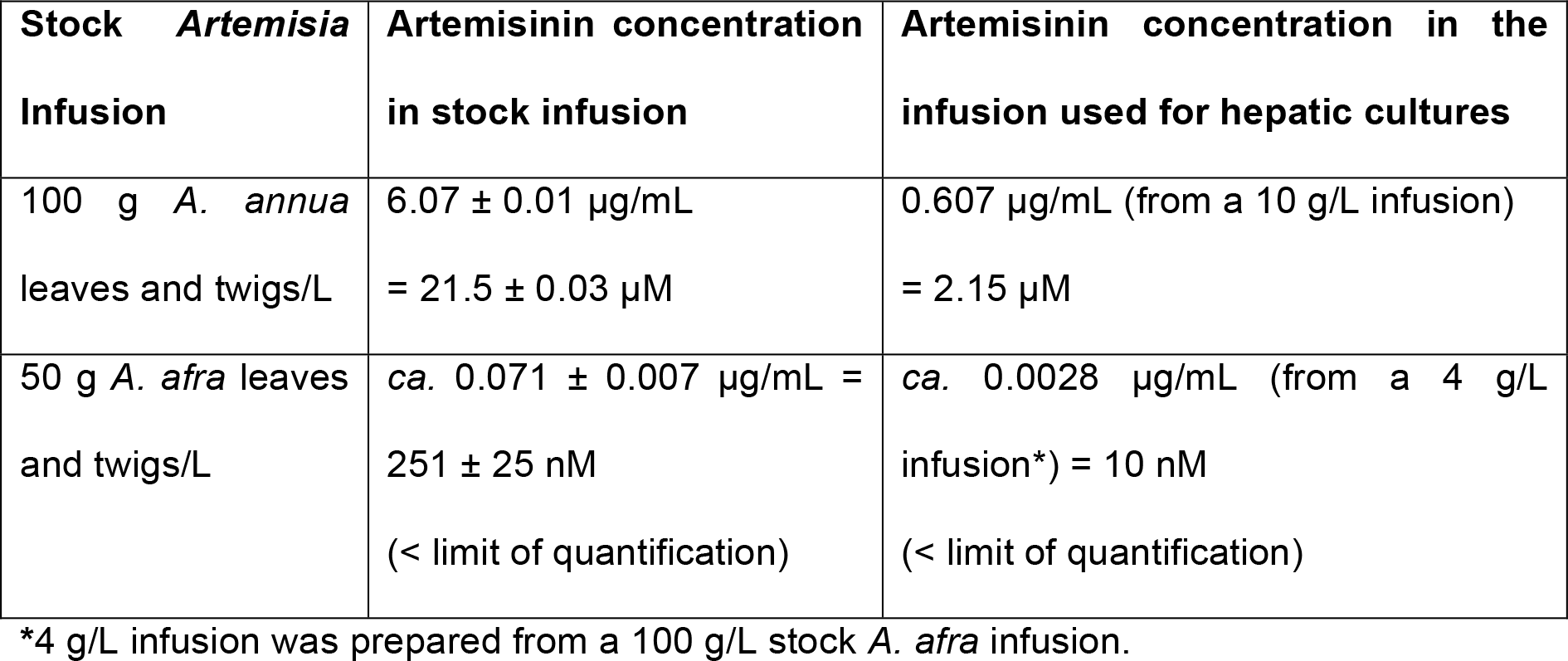
Quantification of artemisinin in *A. annua* and *A. afra* infusion by UHPLC-MS.

### Supplementary Information: Assessment of activity against organellar biogenesis

The inhibitory activity of the infusions is not primarily due to a loss of the 1-deoxy-D-xylulose 5-phosphate (DOXP) pathway (demonstrated to be the only apicoplast function essential for *in vitro* growth of the erythrocytic parasites in lipid-rich medium^1,2^), since similar inhibition by the infusions was observed in apicoplast-negative parasites generated via chloramphenicol (Cm) treatment that were grown in media supplemented or not with isopentenyl pyrophosphate^1,3^ (IPP) **(Supplementary Data Fig. 2a for the study design)**. It also appears that both infusions exhibit a strong growth inhibition at low exposition time (6h), not rescued by the IPP addition **(Supplementary Data Fig. 2b-e)**. The observed apicoplast loss is supported by a rapid loss overtime of the apicoplast genome in infusion-treated cultures, which parallels that of doxycycline-treated positive control cultures^1,4^ **(Supplementary Data Fig. 2f)**. Furthermore, quantification of mitochondrial DNA **(Supplementary Data Fig. 2g)** during the 48-hour *P. berghei* hepatic cycle along with the microscopic imaging **(Fig. 2d)** also revealed disruption of this organelle^1,4^. In both control and treated conditions, parasites continued to augment their DNA and replicated during the 48-hour intracellular development, but this increase in DNA was less obvious in doxycycline and infusion exposed groups **(Supplementary Data Fig. 2h)**.

To further assess whether infusion treatments exert organelle-specific inhibition, synchronized rings were exposed to 0.5 g/L *A. annua* or *A. afra* for 24 h and 48 h and apicoplast development was assessed and compared to those treated or not with 0.8 nM atovaquone (ATQ). Indeed, ATQ is a drug that is currently in use to treat human malaria, and which kills the parasite by targeting its mitochondrion, thus not directly affecting the apicoplast biogenesis. Results at 24 h show a slight but not significant effect of the ATQ treatment on apicoplast biogenesis compared to control or DMSO but both infusion treatments significantly increase the percentage of affected apicoplast with a stronger effect for *annua-*exposed parasites **(Supplementary Data Fig. 3a)**. Apicoplast elongation and branching seemed blocked, unlike ATQ treatment, with unelongated apicoplast after *afra* treatment and apicoplast signal is diffused in the parasite cytosol with vesicles for *annua* **(Supplementary Data Fig. 3a)**. After 48 h of treatment **(Supplementary Data Fig. 3b)**, the percentage of parasite with affected apicoplast was even higher for the two infusions but also for ATQ, which in this case was similar to *afra* treatment. Microscopic imaging of these *P. berghei* hepatic cultures also indicated a disruption of this organelle **(Supplementary Data Fig. 3c)**. Together, this strongly suggests that apicoplast biogenesis might be indirectly affected by the overall death of the parasite after a longer exposure to the different treatments (*afra* or ATQ). But the stronger effect of *afra* at early stage (24 h) indicates a possible direct impact on the apicoplast. Hence, *annua* also highly affect the general development of the parasites, by blocking them at the late ring / early trophozoite stage upon 24 h. However, a high around 60% of the parasites harbour a single punctate apicoplast while ∼40% harbour a disrupted/diffused apicoplast, which could also indicate an impact on the apicoplast but also and meanly a stronger impact on an apicoplast off-target that block the parasite development after short exposition to the infusion (concomitant with Supplementary Data **Fig. 2d**, which shown a drastic negative impact of *annua* after only 6 h of exposure).

**Supplementary Data Figure 1.**
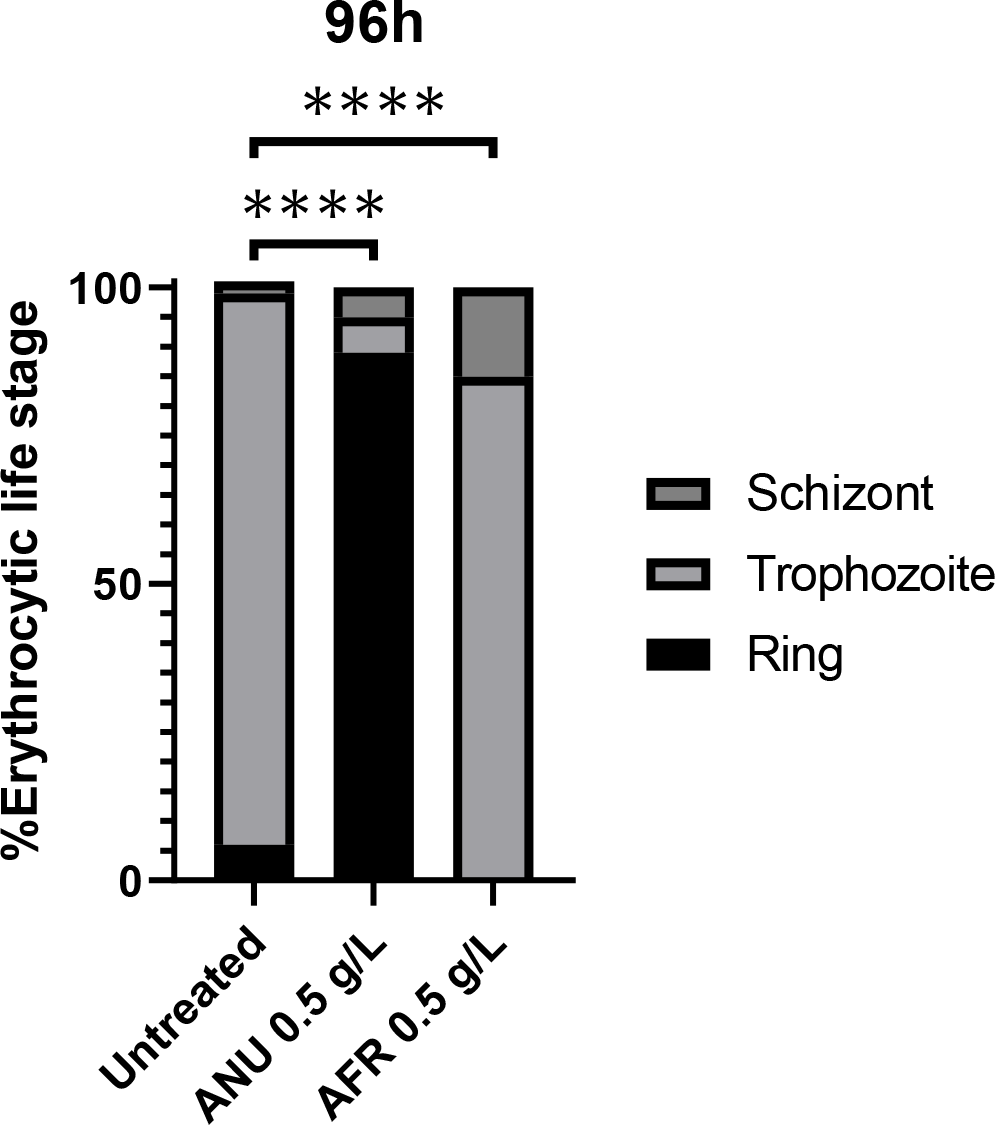
Impact of *A. annua* (ANU) and *A. afra* (AFR) infusions on the development of asexual blood stage of *P. falciparum* from tightly synchronized cultures at ring stage (1% total parasitaemia) followed by two life cycles under infusion treatment (0.5 g/L of infusion) or control (sub-culture at 48 h).

**Supplementary Data Figure 2.**
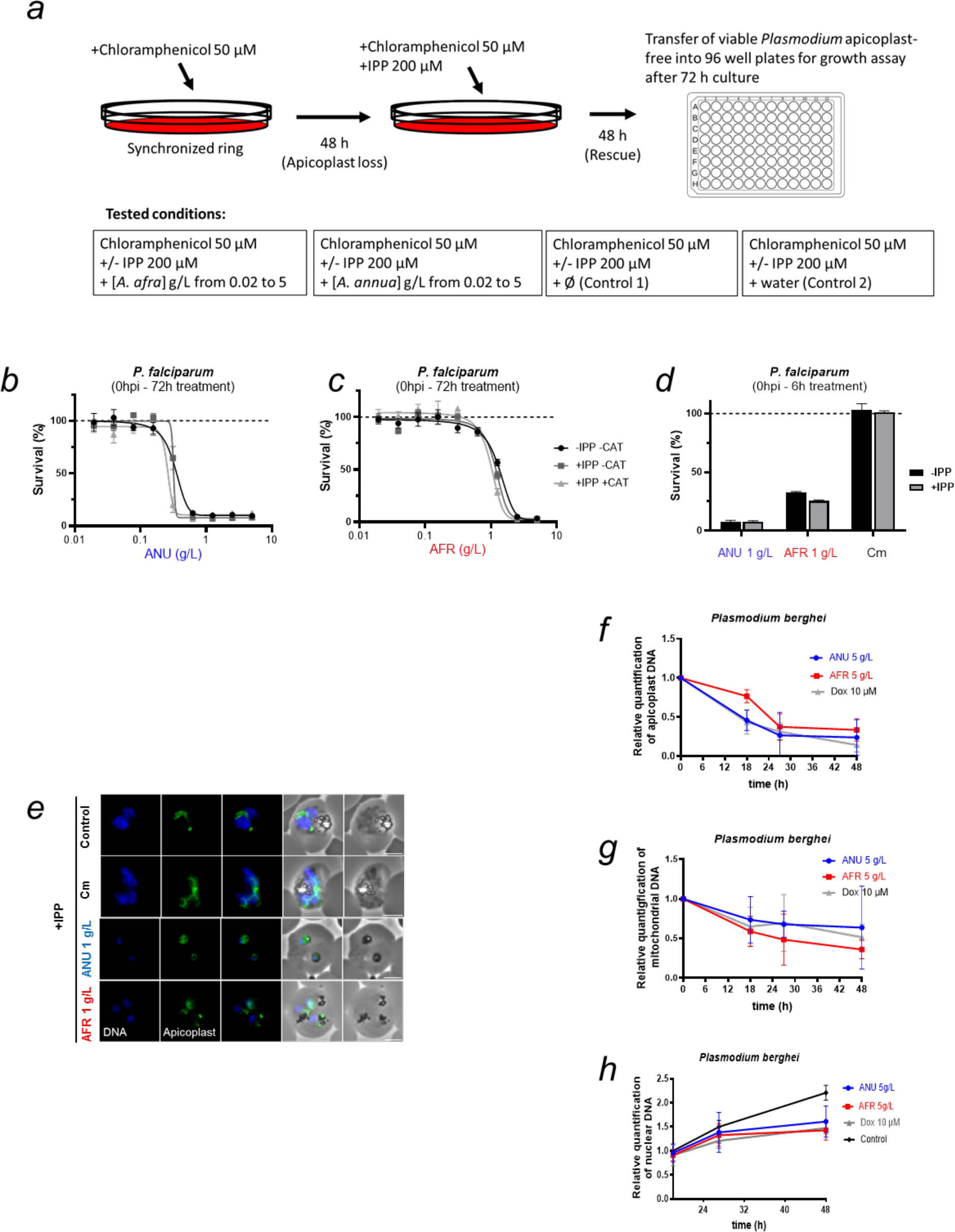
**a,** Schematic of apicoplast loss by chloramphenicol treatment and IPP rescue performed for Fig. 2 b-e. (**b, c**) Parasite survival after 72 h exposure to *A. annua* (**b**) or *A. afra* (**c**) infusions in normal cultures (-IPP/-Cm), apicoplast-free parasites (+Cm) in the presence or absence of IPP. **d, e**, Growth assay and **f**, epifluorescence microscopy of infusion, Cm treated (6 h treatment on ring stage + washes followed by 66 h growth) *P. falciparum* blood stage in the absence or presence of IPP. Scale bar, 3 µm. Results are representative of three (**b** and **c**) or six (**d**) independent experiments. **f, g, h,** Quantification of apicoplast (**f**), mitochondrial (**g**), and nuclear (**h**) genomes by qPCR over the 48-hour development of the *P. berghei* hepatic parasites exposed to infusion or doxycycline. In panels (**f**) and (**g**) each measurement point is normalized to the control of that point and in (**h**) to the control of 18 hours. AFR = *A. afra*, ANU = *A. annua*, ATQ = atovaquone, DOX = doxycycline. Results for **f, g** and **h** are representative of two independent experiments.

**Supplementary Data Figure 3.**
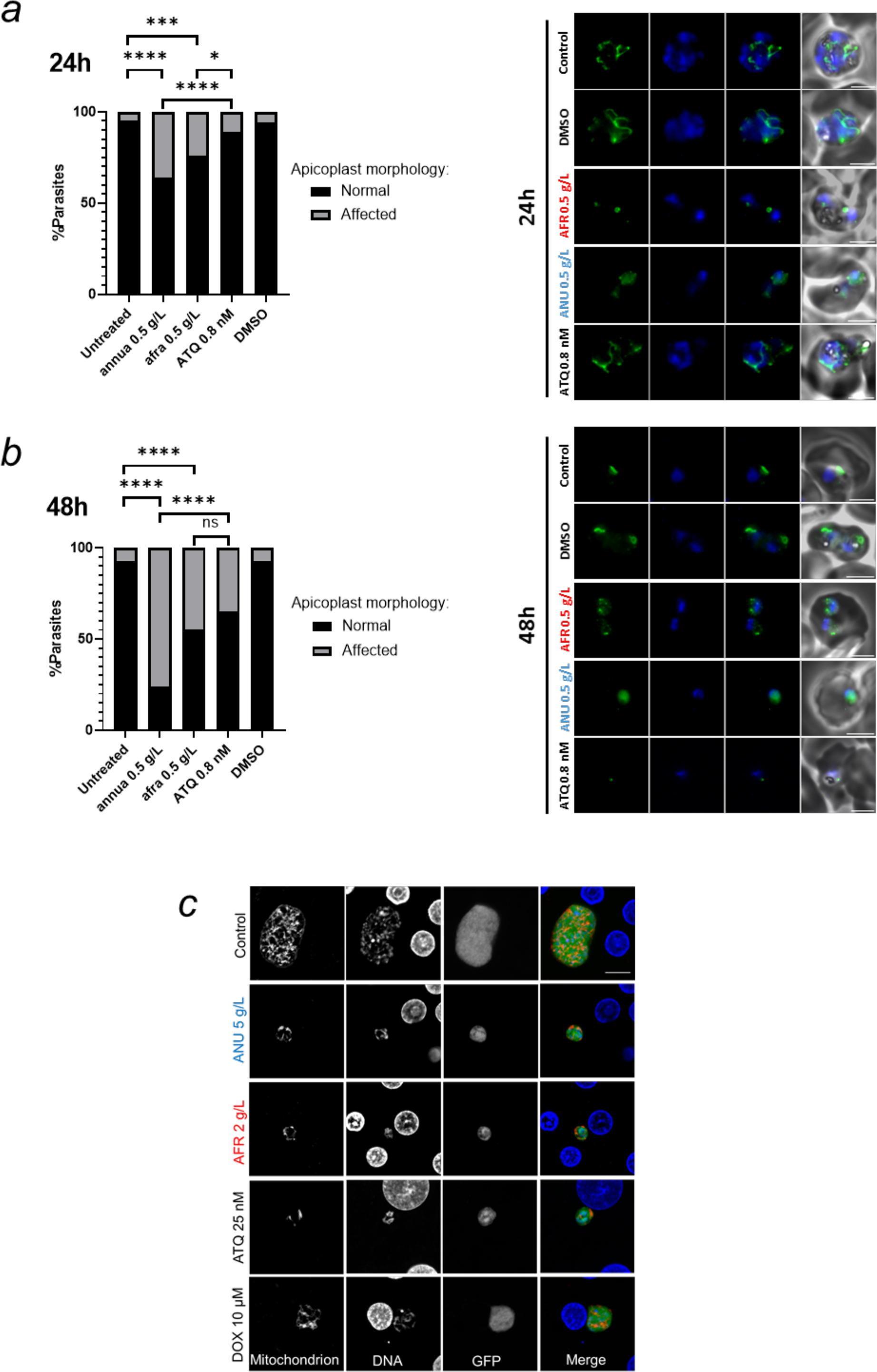
**a, b,** Visualisation of apicoplasts by immunofluorescence in untreated control parasites (trophozoite stage **(a**) and early trophozoite (**b**) or those grown in the presence of the infusions 0.5 g/L or Atovaquone (ATQ) 0.8 nM for 24 h (**a**) or 48 h (**b**) (right panels). Percentage of parasites with apicoplast biogenesis affected by 24 h (**a**) or 48h (**b**) treatments (left panels). **c,** Concentrations of infusions are provided as dry weight of leaves prepared in water and presented as gram per litre (g/L) (See materials and Methods). c, Confocal microscopic images of treated and control *Pb*-GFP mitochondria in infected hepatocytes, as visualized by immunolabeling with anti-recombinant *Pf*HSP60 sera (scale bar = 10 µm).

**Supplementary Data Figure 4.**
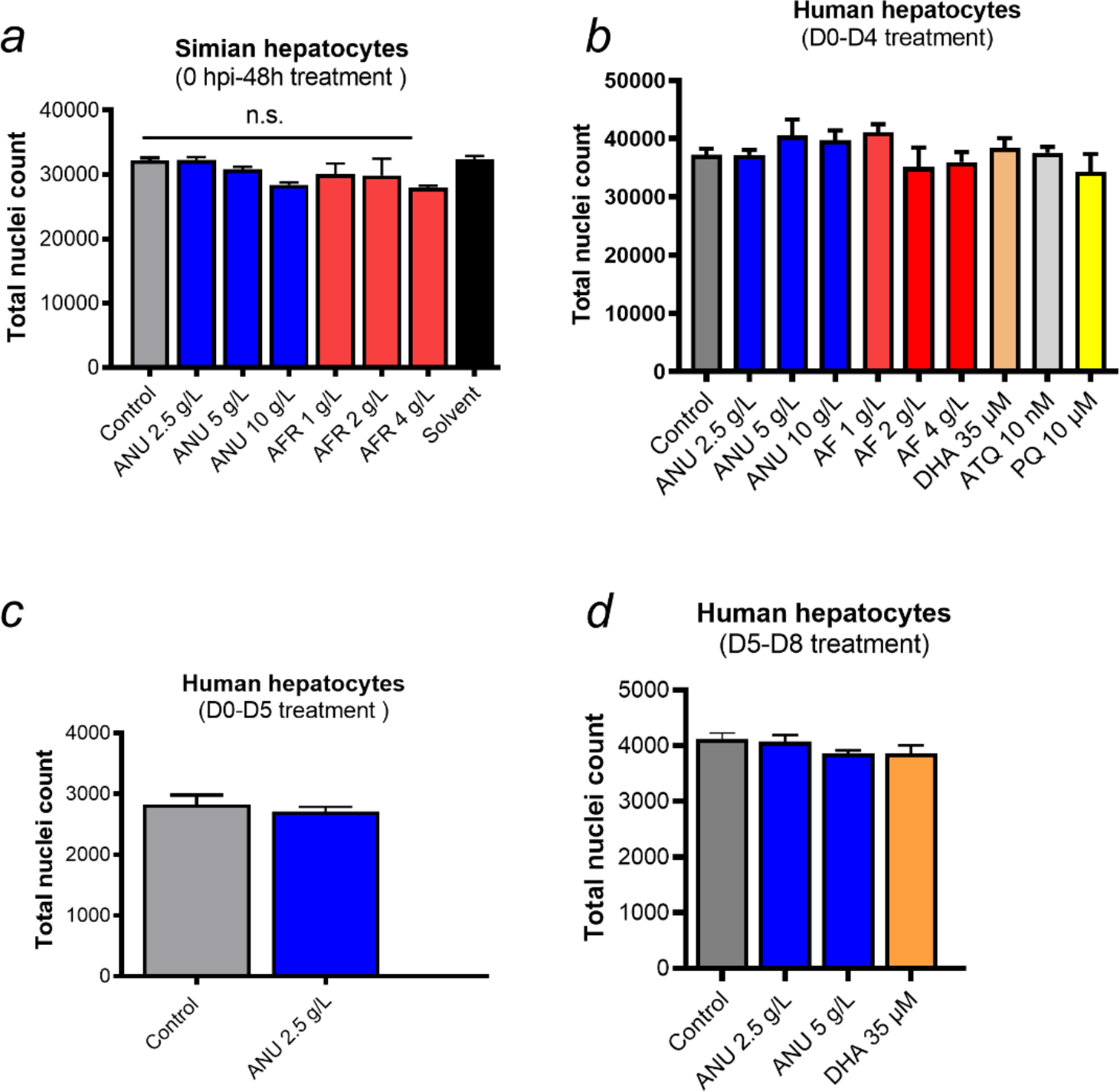
Toxicity of *Artemisia* infusions on simian (**a**) and human (**b-d**) primary hepatocytes. Hepatocyte nuclei are counted from an equal set number of microscopic fields of controls and infusion-treated cultures. Results are representative of three independent experiments.

**Supplementary Data Figure 5.**
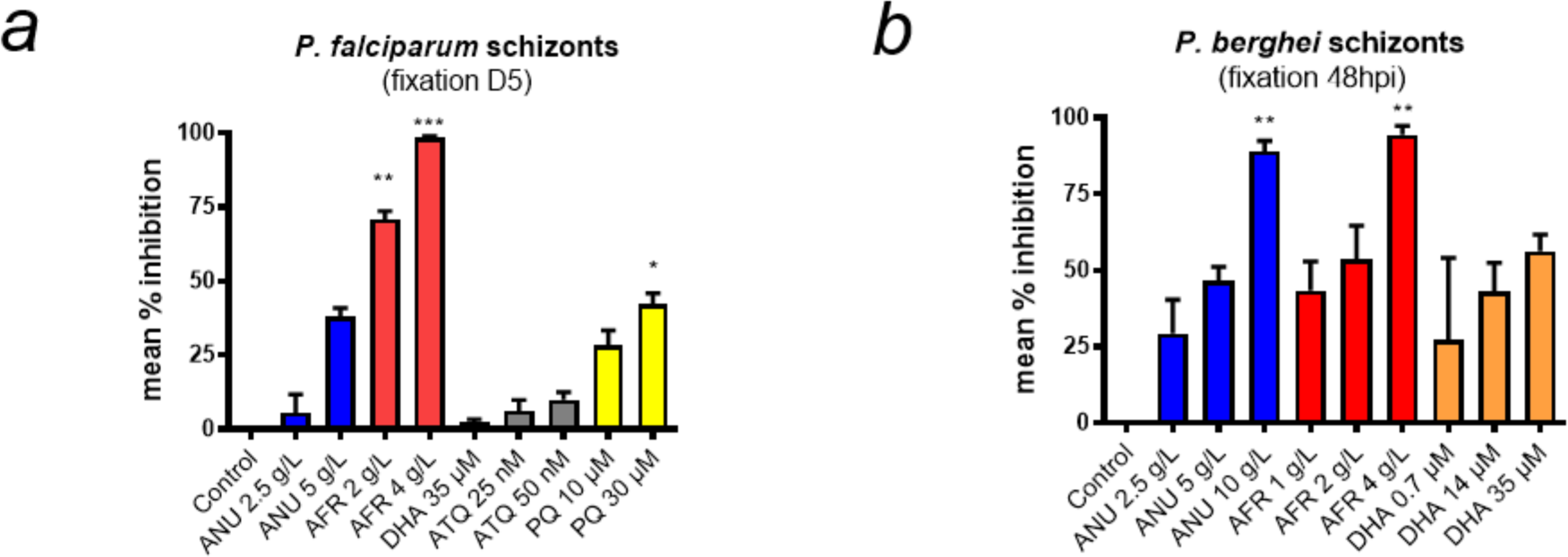
Percent inhibition of *P. falciparum* (**a**) and *P. berghei* (**b**) hepatic schizonts observed in cultures inoculated with sporozoites and incubated with *Artemisia* infusions or drugs for 1 hour prior to addition to the hepatocytes. The exposed sporozoites were washed free of infusion or drug before being allowed to infect hepatocytes. *P. falciparum*-infected human hepatocyte cultures and *P. berghei*-GFP infected simian hepatocytes were maintained until day 7 and day 2 pi, respectively, before parasite enumeration by computational scanning of the plates. *P. falciparum* schizonts were labelled with anti-HSP70 antibody before scanning. The results are representative of three independent experiments. Control wells had on average 552 parasites/well for *P. falciparum* and 185 parasites/well for *P. berghei*. Statistical significance was determined with GraphPad Prism 8 using a one-way ANOVA followed by Kruskal-Wallis Test (multiple comparisons to Control) where significance was evaluated by P < 0.0001 (****), P<0.0013 (**) and P < 0.05 (*).

**Supplementary Data Figure 6.**
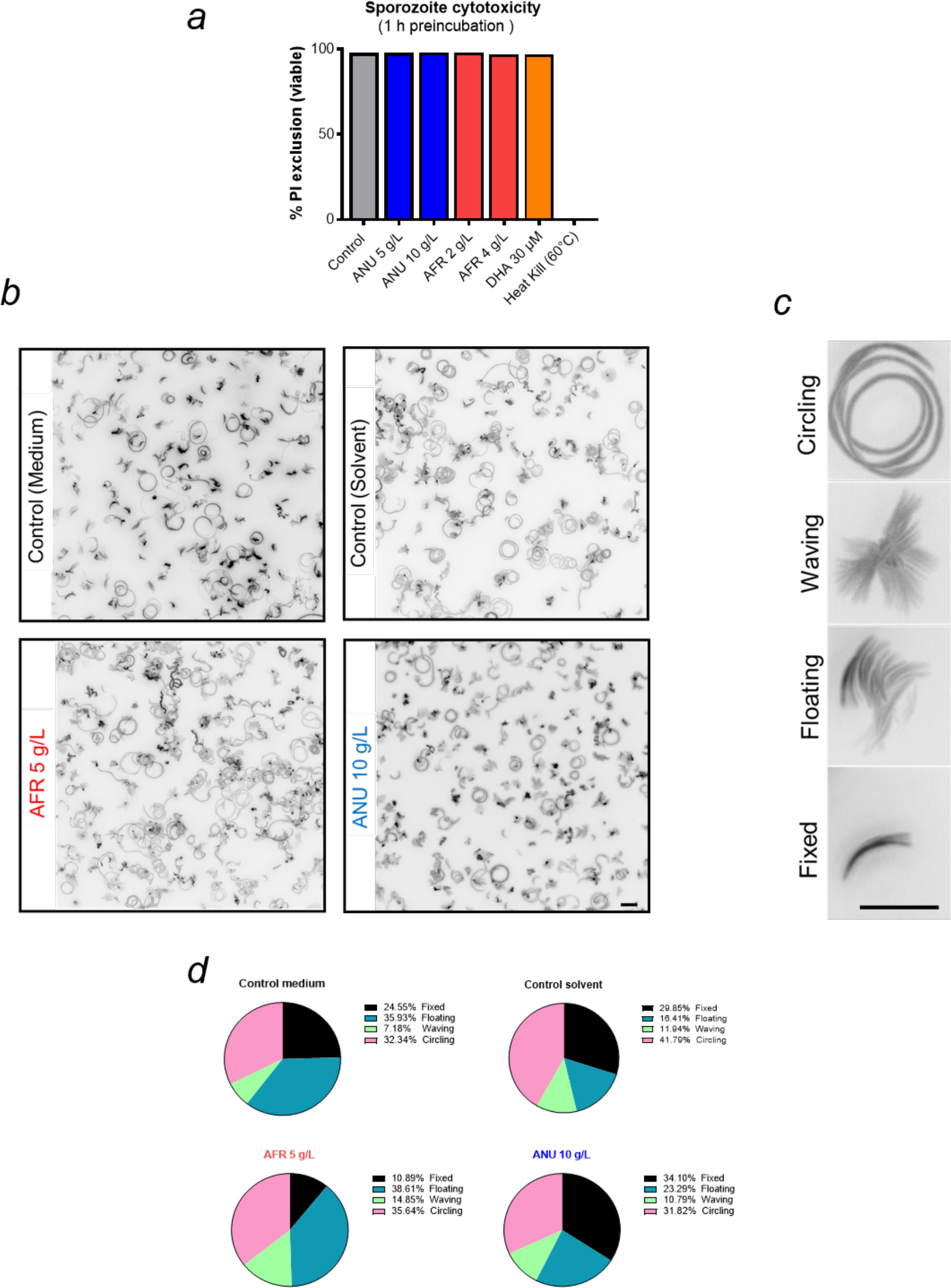
The effect of infusion treatment on free *P. berghei-*GFP sporozoites motility and viability. **a,** Toxicity of infusions on freshly dissected *P. berghei* sporozoites exposed for one hour at room temperature. For each treatment condition, all the movie frames recorded movie were stacked, and a single composite black and white image was prepared using ImageJ software. **b, c, d,** Four different patterns of sporozoite movement (**c**) were identified based on a previous publication^5^ and the frequency of each pattern quantified (**d**). Treatment with infusion did not result in a significant shift from that of controls (scale bar = 10 µm).

### Supplementary Information: Comparative analysis of *A. annua* and *A. afra* secondary metabolites with anti-plasmodial activity and their biological targets

#### Terpenoids

Species of the Asteraceae plant family are rich in anti-plasmodial terpenoids other than artemisinin and congeners, including in those of the *Artemisia* genus.^6–10^ Due to their hydrophobicity, sesquiterpenes are virtually absent in *A. annua* aqueous infusions, with the exception of arteannuin B, dihydroartemisinic acid, and significantly, artemisinin.^11–14^ The terpenoid content of *A. afra* aqueous infusions, to the best of our knowledge, was never investigated except for the absence - or extremely low amount - of artemisinin.^6,15–18^ However, many of the volatile monoterpenes that constitute *Artemisia* essential oils (EO) can be predicted to be partly extractable by warm water, but can only be detected by GC-MS analysis.^19,20^ Among these, 1,8-cineole (*syn*. eucalyptol) and especially artemisia ketone are consistently abundantly found in both *A. annua* and *A. afra* EO.^21–30^ On the other hand, *A. afra* contains a high amount of thujones^21,28,29,31^ and 6,7-epoxylinalool,^32^ which are absent from *A. annua*.^24,25,33^ Minor specific compounds include limonene and linalool in *A. annua*,^33–35^ which are extremely rare or absent in *A. afra*.^28^ While the electrophilic 6,7-epoxylinalool was never tested against malarial parasites, artemisia ketone and thujones were shown to have very weak activity against intra-erythrocytic *P. falciparum* stages,^31^ whereas eucalyptol,^31,36^ linalool^37^ and limonene^37,38^ inhibited parasite growth at elevated (high micromolar to low millimolar) concentrations. Mechanistically, linalool and limonene exert anti-plasmodial effects on the parasite’s blood stages by inhibiting enzymes downstream of the apicoplast DOXP/MEP pathway (i. e., prenyl synthases and protein prenyltransferases), presumably by competing with their isoprenoid substrates.^37,38^ Stronger effects can be found in related non-*Artemisia* monoterpenes (e.g., perillyl alcohol) and sesquiterpenes (e. g., nerolidol). These inhibitors repress the biosynthesis of dolichols and prenylated proteins,^37–40^ as well as that of the octaprenyl chain of ubiquinone,^41^ thus potentially affecting the mitochondrion.^42^ In the case of *A. annua*, the presence of significant amount of artemisinin could partly contribute to a direct disruption of the parasite mitochondria.^43–45^ Interestingly, no natural terpenoid has yet been found to inhibit the DOXP/MEP pathway in the apicoplast as does fosmidomycin.^46^ Due to the existence of established or putative anti-plasmodial terpenes in the rich EO of *A. annua* and *A. afra*, it appears important in the future to measure their quantity in *Artemisia* infusions, and also to test representative compounds (e. g., eucalyptol) against the parasite liver stages with a focus on anti-apicoplast and anti-mitochondrion effects.

#### Flavonoids and chlorogenic acids

Aqueous infusions of *A. annua* contain a major proportion of polar metabolites dominated by flavonoid aglycones (e. g., jaceidin, casticin), flavonoid glycosides (e. g., apigenin-6-*C*-glucopyranoside, *syn*. *iso*-vitexin, luteolin-7-*O*-glucopyranoside, *syn*. cynaroside) and quinic acid esters (*syn*. chlorogenic acids).^11,12,23,47–49^ *A. afra* aqueous extracts were never characterized in-depth except for the qualitative detection of flavonoids and phenolics.^50^ However, this species is known to produce flavonoid glycosides common to *A. annua* (e. g., cynaroside) in addition to several flavonoid aglycones (e. g., apigenin, tamarixetin)^6,51,52^ which albeit distinct from those present in *A. annua* can be predicted to parallel or even surpass their hydrosolubility. *A. afra* also contains an important component of chlorogenic acids.^17,53^ These *A. afra* metabolites could thus be present in aqueous infusions of A. afra, similarly to those of *A. annua*. Most flavonoids and chlorogenic acids from both *Artemisia* species exert significant inhibition on blood-stages *Plasmodium* parasites, with activities ranging between low and high micromolar.^6,11,17,47,51,54–59^ The bulk contribution of these metabolites to the activity of *A. annua* infusions against *P. falciparum* erythrocytic stages, which is dominated by a both abundant and extremely potent artemisinin, was shown, however, to be negligible using plant strains deficient in either flavanone (Δ*CHI1-1*) or amorpha-4,11-diene (*Si*AMS or Δ*cyp71av1-1*) biosynthetic pathways.^60,61^ On the other hand, artemisinin activity falls by *ca*. three orders of magnitude against the parasite liver stages,^62,63^ consistent with their fewer endoperoxide-activating heme sources compared to those in actively-digesting blood parasites.^45,64–66^ In this context, most of the aforementioned compounds present in *A. annua* or *A. afra* - the latter species notably lacking artemisinin - may start to exhibit significant inhibition of *Plasmodium* hepatic stages. Mechanistically, certain anti-plasmodial flavonoids present in *A. annua* infusion (i. e., cynaroside) or abundant in *A. afra* (i. e., apigenin) were shown to inhibit the *P. falciparum* enoyl-acyl carrier protein reductase (PfENR, *syn*. FabI),^57,58^ an enzyme harboured by the parasite type II fatty acid synthase (FASII) machinery in the apicoplast.^46^ Other flavonoids from aqueous extracts of both *Artemisia sp.* could also be found to target the apicoplast, considering the low micromolar activity of various congeners against FabG, FabZ or FabI,^58^ all part of plasmodial FASII complex.^46^ Regarding the anti-mitochondrial effects, many of the anti-plasmodial flavonoids present in *A. annua* infusions (i. e., *iso*-vitexin, vitexin and casticin) or present *in A. afra* (e. g., apigenin, tamarixetin) are known to disrupt the inner mitochondrial membrane potential, to generate reactive oxygen species (ROS) and to induce apoptosis in cancer cells.^67–79^ In plants, *iso*-vitexin was shown to disrupt mitochondrial respiratory chain by interacting with ubiquinone binding sites.^80^ In *Leishmania* parasites, apigenin was responsible for ROS generation, mitochondria depolarization and swelling.^81^ Similar mitochondrion-targeting mechanisms could underpin the action of these flavonoids in *Plasmodium* parasites, although this has never been demonstrated. On the other hand, certain anti-plasmodial flavonoids from both *Artemisia* species (i. e., cynaroside, chrysoeriol) possess ROS-scavenging capability, with mitoprotective and antiapoptotic effects,^82–84^ leading to tentatively exclude mitochondrion-disrupting mechanisms for these compounds in the parasite. The molecular targets of chlorogenic acids in *Plasmodium* are unknown, but the fact that chlorogenic acid itself was predicted to bind FabI *in silico*^85^ validates their investigation for anti-apicoplast effects related to FASII inhibition. Regarding possible mitochondrial effects, chlorogenic acid and congeners show cytotoxicity and induce mitochondrion-dependent apoptosis in cancer cells *via* ROS production and collapse of the inner membrane potential, in addition to other effects (e. g., inhibition of the PI3K/AKT/mTOR pathway, activation of PKC).^86–90^ Interestingly, these effects seem to be inexistent in normal cells, where chlorogenic acid exerts SIRT1-dependent antioxidant, mitoprotective and antiapoptotic activities.^88,91–94^ Hence, similarly to *Artemisia* flavonoids, the possibility of beneficial or detrimental effects of chlorogenic acids makes it difficult to predict their action in *Plasmodium* mitochondria.

The artemisinin-independent anti-plasmodial effects exerted by *Artemisia annua* and *A. afra* infusions in the present study could well be explained by a concomitant, multi-target action of various moderately potent phytochemicals against *Plasmodium* parasites. However, the existence of novel, yet undetected minor compound(s) endowed with hyperactivity, constitutes a hypothesis worth being investigated in a drug discovery perspective.

